# Environmental RNA Improves Detection and Surveillance of Schistosomiasis Transmission

**DOI:** 10.64898/2026.01.19.699857

**Authors:** Chiara Mercier, Philippe Douchet, Damien Pouzol, Jean-François Allienne, Roberta Lima Caldeira, Marina Moraes Mourão, Mariana Gomes Lima, Lângia Colli Montresor, Simon Blanchet, Géraldine Loot, Olivier Rey

## Abstract

Environmental diagnosis of schistosomiasis, a tropical disease affecting more than 250 million people globally, is still challenging, which limits efficient eradication plans. There is a crucial need for resolutive and highly sensitive environmental tools to improve disease control. However, a major obstacle is the inability of current methods, including environmental DNA (eDNA), to distinguish infectious parasite life stages. Here, we develop and validate an environmental RNA (eRNA) approach that enables the detection and absolute quantification of *Schistosoma mansoni* miracidia and cercariae directly from water samples. After identifying life stage–specific transcripts, we design specific ddPCR assays which are strongly specific to *S. mansoni* and to each life stage. Comparing with eDNA assays, laboratory experiments demonstrate that stage-specific eRNA assays accurately reflect the relative abundance of each life stage, detect nucleic acids released from organisms, exhibit detection limits tenfold lower than eDNA assays. Also, under laboratory conditions, RNA persists in water longer than DNA. Field validation at six endemic sites in Brazil confirms that eRNA outperforms eDNA and conventional snail surveys in detecting active presence of *S. mansoni* cercariae, which indicates schistosomiasis transmission risk to humans locally. By enabling active-stage discrimination in environmental monitoring and improving sensitivity (compared to eDNA), our study advances both fundamental understanding and applied surveillance of schistosomiasis transmission, supporting elimination initiatives in affected regions.

## INTRODUCTION

Schistosomiasis or bilharzia is a disease caused by parasitic blood-dwelling flukes of the genus *Schistosoma* that persists as a neglected tropical disease (NTD) imposing a significant burden on public health (1, 2). Over 250 million people are infected across 78 countries, with nearly 800 million people at risk, predominantly in tropical and subtropical regions where access to safe water and adequate sanitation remains limited (3–5). Despite a global decline in prevalence in recent years, the general persistence of transmission underscores the need for effective tools enabling early detection of transmission foci and improved disease control.

Control and elimination strategies have traditionally emphasized praziquantel-based mass drug administration to humans, expanded surveillance, and improved access to clean and safe water (6). However, ongoing transmission remains challenging to interrupt due to the parasite’s complex life cycle (Fig. 1). Schistosomes alternate between long lifespan within their definitive vertebrate hosts (humans or animals) and specific freshwater snails, which serve as intermediate hosts. Following release into the aquatic environment with definitive hosts’ feces or urine, parasite eggs hatch into miracidia, which actively seek and infect compatible snail hosts. Asexual development within the snail produces hundreds to thousands of infective clonal cercariae per day, that are released into the water, where they penetrate the skin of the definitive host to complete the cycle (1, 3). Therefore, local and seasonal transmission dynamics of schistosomiasis are intrinsically linked to freshwater ecosystems, spatial and temporal heterogeneities in the distributions of snail host, potential vertebrate reservoir host, and to human water-use behaviors (7–9).

**Fig. 1.**
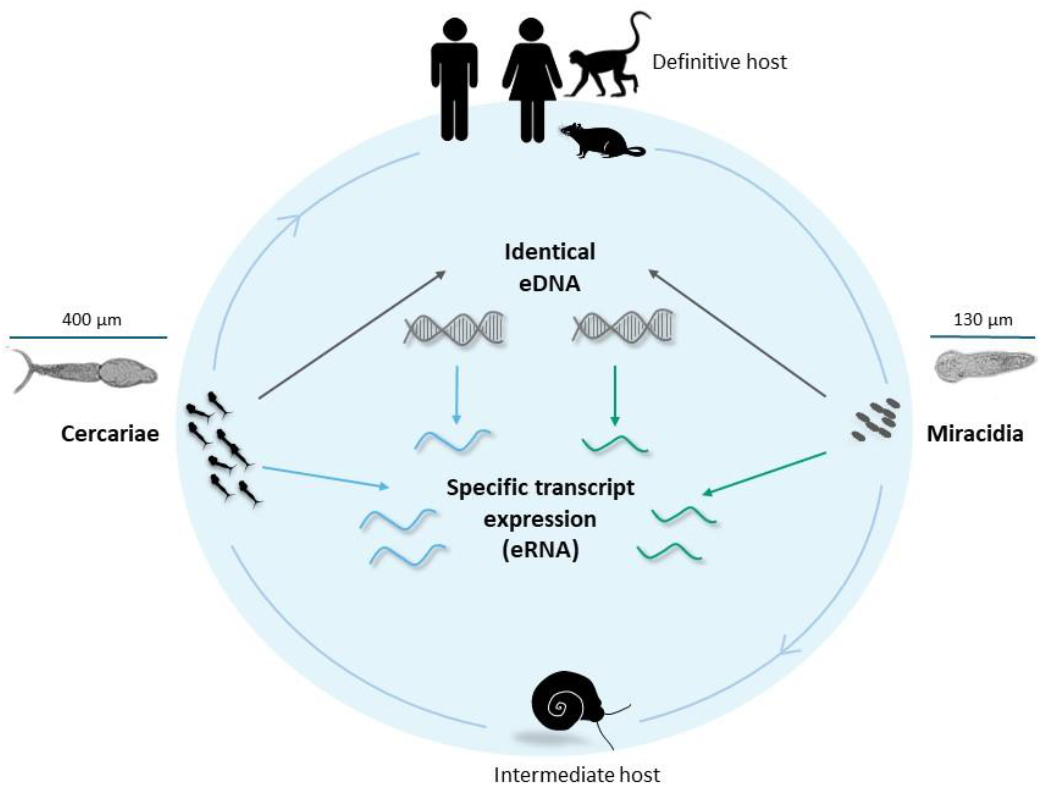
Representation of the life cycle of *Schistosoma mansoni* with a focus on active infectious life-stages and their expected environmental nucleic acid signals. Eggs released by the definitive host (human or animal) hatch into miracidia (∼130 µm) which are infectious to snails. Inside the snail, the miracidia develop in sporocyst which multiply asexually. Then the cercariae (∼400 µm) are emitted by the snail into the water and infect definitive hosts. Both free-living stages, miracidia and cercariae, release identical environmental DNA into the water. In contrast, they release stage-specific transcript as environmental RNA (eRNA), which may provide a precise signal of the parasite stages that are actively infectious.

Monitoring ongoing schistosomiasis transmission in water ecosystems is then essential for evaluating control interventions and preventing resurgence. Traditional surveillance approaches, which include parasitological diagnosis in humans (generally neglecting animal hosts in the case of zoonotic *Schistosoma* species) and malacological surveys targeting infected snails, face major challenges of sensitivity, safety, and scalability, particularly in low-prevalence or intermittent transmission areas (10, 11). The recent discovery that organisms leave DNA imprints in their surrounding environment has revolutionized the surveillance of pathogens, including schistosomiasis. Environmental DNA (eDNA) now provides an alternative tool, enabling direct detection of different schistosome species from water samples with higher sensitivity and spatial resolution than traditional parasitological surveys (11–14). For instance, eDNA-based approaches have successfully identified transmission sites in Africa, South America, and Asia (11, 15–17). Nonetheless, a major drawback of eDNA is its inability to differentiate parasite life-stages (*i*.*e*. miracidia vs. cercariae), since both carry the same genetic information. Moreover, eDNA can persist over time and be transported across the aquatic environment, making it difficult to distinguish between DNA from active infectious parasites and dead or residual parasite sources (18, 19). All these existing surveillance methods may lead to false positive detection, hence limiting an accurate assessment and mapping of the true risks of transmission to humans (12).

Environmental RNA (eRNA) has recently emerged as a promising tool to overcome these limitations (20). Because eRNA reflects gene expression, life-stage specific transcriptomic signature may allow the discrimination among parasite life-stages from environmental samples. For *S. mansoni*, distinguishing miracidia-derived signals (reflecting water contamination by infected definitive hosts) from cercariae-derived signals (indicating direct human infection risk) adds crucial biological and epidemiological resolution to environmental surveillance (Fig. 1). Moreover, because eRNA is produced only by active cells and is generally assumed to degrade rapidly (i.e., has a short temporal persistence and high degradation rate), it can serve as an indicator of active and ongoing transmission (20). Noteworthily, a few recent studies show that RNA molecules can actually persist in the environment longer than expected (>30 days), sometimes even longer than DNA molecules (21, 22). Hence, a longer stability of eRNA may improve detection sensitivity and field applicability, thereby bridging critical gaps in understanding emergence patterns.

Here, we develop and validate an environmental RNA-based assay that detects, discriminates, and quantifies each of the two free-living stages, miracidia and cercariae, of *Schistosoma mansoni* - the major etiological agent of intestinal and hepatic schistosomiasis in Africa and South America - directly from environmental water samples. We evaluate the specificity and sensitivity of stage-specific eRNA digital droplet PCR (ddPCR) assays and compare their performance to an eDNA assay targeting *S. mansoni*. We next conduct three controlled experiments to validate the quantification efficiency of eRNA assays. We test the ability of stage-specific eRNA assays to adequately quantify their respective eRNA target (miracidia or cercariae) across varying proportions of each life-stage (Hypothesis 1), expecting stage-specific RNA signals to reflect the life-stage abundance, while eDNA remains stable across conditions. We also test whether nucleic acids released into the water by organisms (free RNA and DNA molecules) are detectable after organisms are removed from the environment (Hypothesis 2). We anticipate detectable but reduced eDNA and stage-specific eRNA signals than when organisms are still present in the environment (15). Third, we quantify the persistence and decay kinetics of eRNA and eDNA signals over time (Hypothesis 3), expecting RNA to persist at least as long as eDNA. Finally, we apply the stage-specific eRNA assays to endemic field sites in Brazil to compare stage-specific eRNA detection outcomes with eDNA and classical parasitological snail surveys. By integrating eRNA detection into environmental surveillance, this work enables real-time and stage-specific monitoring of transmission dynamics, offering a powerful tool to inform and optimize targeted interventions for local intestinal schistosomiasis control.

## RESULTS

### Stage-specific assays design

Among the genes screened using the ‘SchistoXYZ’ web platform (23), two candidate transcripts were selected for developing stage-specific eRNA markers based on their expression levels across different *S. mansoni* life stages and their optimal expression levels in terms of detectability (SI Appendix, Fig S1). The *Smp_032670* gene, encoding an egg protein C122-like, displayed the highest expression level in miracidia among all genes and was not expressed in cercariae (although being slightly expressed in eggs and sporocysts, a developmental stage in mollusks prior to the cercaria stage). The *Smp_169190* gene, which encodes a tegumental allergen-like protein, displayed the highest expression level in cercariae among the 10 genes initially targeted and no expression in miracidia (SI Appendix, Fig S1). The designed stage-specific eRNA primers targeting these two gene transcripts (Table 1) were validated *in silico* based on the *ncbi* nr database using Primer-Blast. Only *S. mansoni* species was targeted by the primers designed for miracidia. The primers designed for cercariae were predicted to amplify *S. mansoni* and *S. rodhaini* based on the nt database, and *S. mansoni* and *S. haematobium* (with one mismatch per primer) based on the RefSeq RNA database (SI Appendix, Fig S2). Primers and probes were next validated *in vitro* using DNA/RNA extracts from freshly hatched miracidia and cercariae (see next paragraph).

**Table 1.**
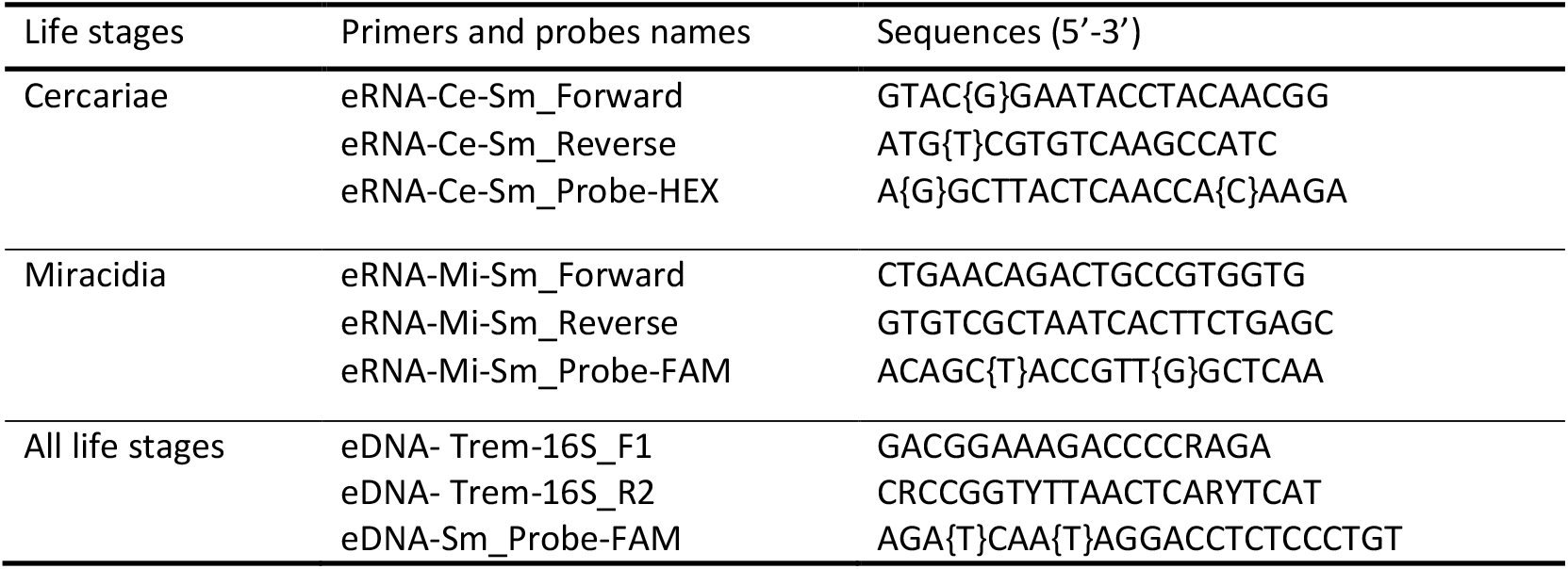
RNA- and DNA-based primers and probes used in this study. The brackets indicate a locked nucleic acid (LNA) that allows enhancing melting temperature (Tm) during PCR amplification.

### Validation of the specificity and sensitivity of the designed assays

Regarding the specificity of the stage-specific eRNA assays, we respectively ran ddPCR on triplicate samples of *S. mansoni* miracidia and cercariae, and mock communities composed of RNA from other trematode species without the presence of *S. mansoni* respective life-stage. The mean concentration of RNA in mock communities was 0 for miracidia and 3.80 copies/µL for cercariae (Table 2), with consistent values across replicates. This low-level signal observed for the cercariae-specific assay indicated existing -but limited-cross-amplification with RNA from other cercariae species. The specificity of the assays was strong among stages (Table 2), since the miracidia-specific assay did not amplify RNA from cercariae (0 copies/µL), and the one developed for cercariae only amplified a minimal RNA signal from miracidia (0.15 copies/µL, compared to 42.56 copies/µL when the cercariae-specific assay was used, see Table 2).

**Table 2.** RNA concentration (copies/µL) of cercariae and miracidia obtained by ddPCR in each tested mock community. Values are expressed as mean ± standard error of the mean (SEM).

| Assay | Mock community or sample tested | RNA concentration of detected target (copies/μL, mean ± SEM) |
| --- | --- | --- |
| **Cercariae assay** | RNA mock without *S. mansoni* cercariae | 3.80 ± 1.11 |
|  | cercariae from *S. mansoni* | 42.56 ± 4.36 |
|  | miracidia from *S. mansoni* | 0.15 ± 0.09 |
| **Miracidia assay** | RNA mock without *S. mansoni* miracidia | 0 |
|  | miracidia from *S. mansoni* | 183.33 ± 9.21 |
|  | cercariae from *S. mansoni* | 0 |

The limits of detection (LOD) of the stage-specific eRNA assays were 0.01 equivalent individual for miracidia and 0.0001 equivalent individual for cercariae (SI Appendix, Table S1). The *S. mansoni* eDNA assay also showed high species specificity, with no amplification from other trematodes, and a LOD of 0.01 equivalent individual for both stages (SI Appendix, Text S1). These results confirm the high species and stage specificity of both stage-specific eRNA assays as well as their high sensitivity.

### Hypothesis 1 - Stage-specific eRNA reliably quantifies *S. mansoni* life stages abundance

To evaluate whether stage-specific eRNA assays reliably quantify the relative abundance of *S. mansoni* life stages, we analyzed water from triplicated experimental tanks containing one of three miracidia:cercariae ratios (5:45, 25:25, 45:5). Stage-specific eRNA and total *S. mansoni* eDNA were quantified using ddPCR. As expected, eRNA concentrations for each *S. mansoni* life stage in tanks with unbalanced ratio (*i*.*e*. 5:45 and 45:5) did not derived from the expected concentrations computed based on raw quantification obtained at the equilibrated the 25:25 ratio (Fig. 2). A Generalized Linear Model (GLM) revealed a significant interaction term between the nucleic acid assay type, and the miracidia:cercariae ratio treatments (SI Appendix, Table S2). This interaction term indicated that each assay (stage-specific eRNA or eDNA) varies differently according to miracidia:cercariae ratio treatments. Indeed, the eDNA assay revealed similar DNA concentrations across the three ratios (Figure 2, SI Appendix, Table S3. b), whereas RNA concentrations revealed by stage-specific eRNA assays varied significantly across ratios (Figure 2, SI Appendix, Table S3. a). Specifically, the miracidia-specific eRNA assays yielded significantly higher RNA concentrations in the miracidia-biased treatment (cercariae RNA *vs*. miracidia RNA: z = -3.58, p = 0.001; SI Appendix, Table S3. a), whereas the cercariae-specific eRNA assays revealed higher RNA concentrations in the cercaria-biased treatment (cercariae RNA *vs*. miracidia RNA: z = 13.19, p < 0.0001, SI Appendix, Table S3. a). At the balanced 25:25 ratio, eRNA copy numbers for each life stage were consistently higher than DNA copy numbers, despite eDNA representing the combined contribution of both stages (DNA *vs*. cercariae RNA: z = -11.77, p < 0.001; DNA *vs*. miracidia RNA: z = -6.78, p = 0.001) (Figure 2, SI Appendix, Table S3. b). This pattern holds true across all treatments with varying miracidia:cercariae ratios (Figure 2). In addition, cercariae-derived RNA concentration was systematically much higher than miracidia-derived RNA at a balanced ratio (z = 5.06, p < 0.001). These results demonstrate that stage-specific eRNA assays accurately reflect the relative abundance of *S. mansoni* life stages in mixed environmental samples, whereas eDNA remains insensitive to changes in life-stage composition.

**Fig. 2.**
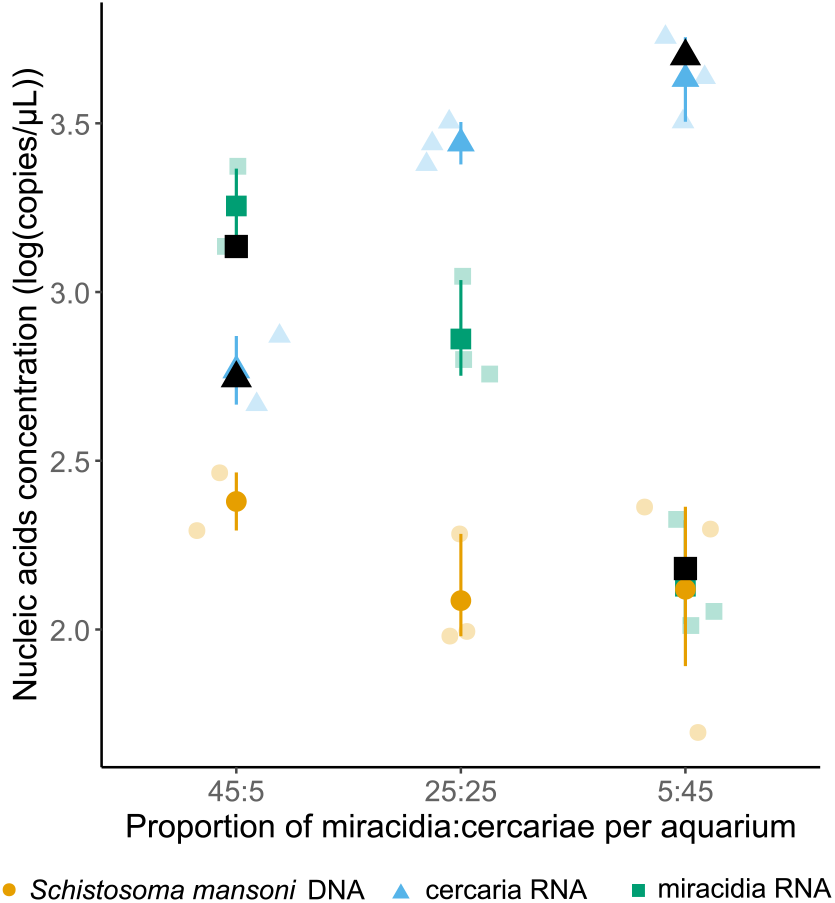
Concentration in log(copies/µL) of *Schistosoma mansoni* DNA (orange circle), cercariae RNA (blue triangle), and miracidia RNA (green square) under three experimental proportions of miracidia:cercariae conditions (45:5, 25:25, 5:45). Semi-transparent points represent observed values from each replicate, filled points indicate the mean of the replicates. Black symbols show the expected eRNA concentrations of miracidia and cercariae in the water based on the mean eRNA concentrations for each life stage obtained from the 25:25 ratio treatment. Negative-control tanks consistently remained free of detectable *S. mansoni* DNA and RNA

### Hypothesis 2 - Released free eRNA and eDNA are detectable

To assess the ability of the eRNA and eDNA assays to detect free nucleic acids released into the water (without the organisms), we used experimental tanks containing a balanced miracidia:cercariae ratio (25:25). Nucleic acid concentrations were compared between filtered water from tanks in which *S. mansoni* life-stages were removed (only free nucleic acids present) or not (both free nucleic acids and organisms present) 5 h post-exposure. eDNA and stage-specific eRNA were both detectable in the water from “free nucleic acids-only” tanks (Fig. 3). However, for all three nucleic acid types (*i*.*e*. miracidia RNA, cercariae RNA, and *S. mansoni* DNA), the removal of organisms significantly reduced the average detected copy number by 1.46, 2.32, and 1.08 log(copies/μL), respectively (likelihood ratio test: χ^2^, p < 0.001, SI Appendix, Table S4). These results demonstrate that free nucleic acids released by miracidia and cercariae can be detected using the eRNA and eDNA assays, supporting their applicability in the field, even in the absence of organisms.

**Fig. 3.**
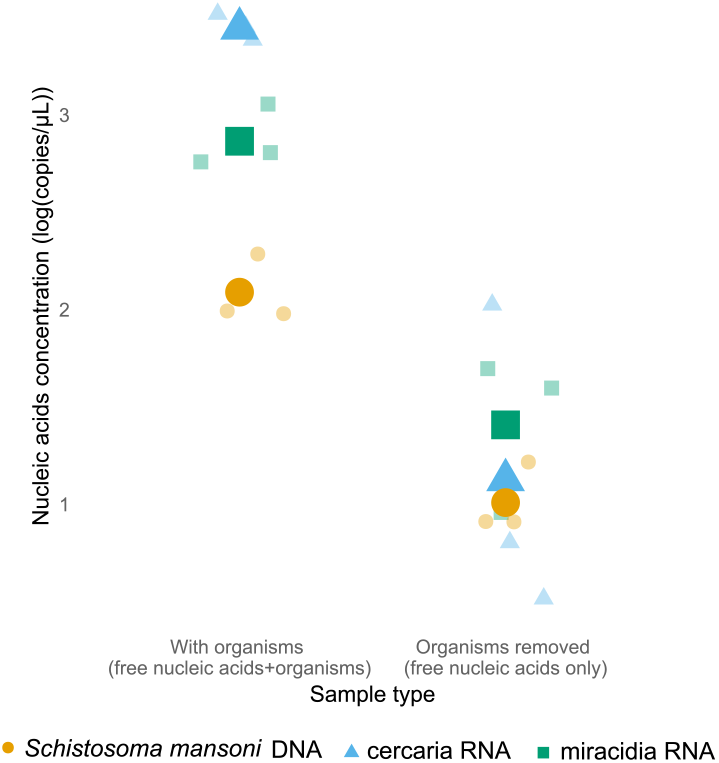
Concentrations in log(copies/µL) of *Schistosoma mansoni* DNA (orange circle), cercariae RNA (blue triangle), and miracidia RNA (green square) in water containing whole organisms (miracidia and cercariae) (with organisms) and in water prefiltered using a 25µM mesh net to discard whole organisms (organisms removed). Semi-transparent points represent observed values from each replicate, filled points indicate the mean among replicates. Negative-control tanks consistently remained free of detectable *S. mansoni* DNA and RNA

### Hypothesis 3 - eRNA persists as long as the eDNA signal in experimental tanks

To characterize the persistence and decay of *S. mansoni* eDNA and stage-specific eRNA signals, we conducted a time-series tank experiment with sampling across multiple days. On day 0, we set up 15 waters tanks containing equal proportions of miracidia and cercariae. At each planned filtration time point (days 0, 1, 2, 7, and 12) three replicates were filtered simultaneously. All nucleic acids were quantified using ddPCRs using both the eDNA and stage specific eRNA assays. A GLM revealed a significant interaction term between the nucleic acid assay type and the sampling day, indicating distinct degradation dynamics for miracidia-derived eRNA, cercariae-derived eRNA, and eDNA (SI Appendix, Table S5). While the concentration of detected miracidia-specific RNA declined rapidly from the first day onward and became undetectable after day 2, both eDNA and cercariae-specific eRNA concentrations remained high after 24 hours and only dropped between days 2 and day 7 (SI Appendix, Table S6). Cercariae-specific eRNA concentrations were significantly higher than those of the other nucleic acids, regardless of the sampling day, and remained detectable at day 7 (whereas eDNA was not detectable) and somewhat detectable (although inconstantly across replicates) at day 12 (Fig. 4). Variability in the detection ability among replicates increased over time, consistent with stochastic degradation processes at low concentration (Fig. 4). The observed decay dynamics demonstrate that eRNA – more precisely cercaria-specific eRNA - in addition to being more biologically informative, tended to persist longer than eDNA (at least in some replicates), suggesting that it could be a more sensitive approach for *S. mansoni* environmental detection.

**Fig. 4.**
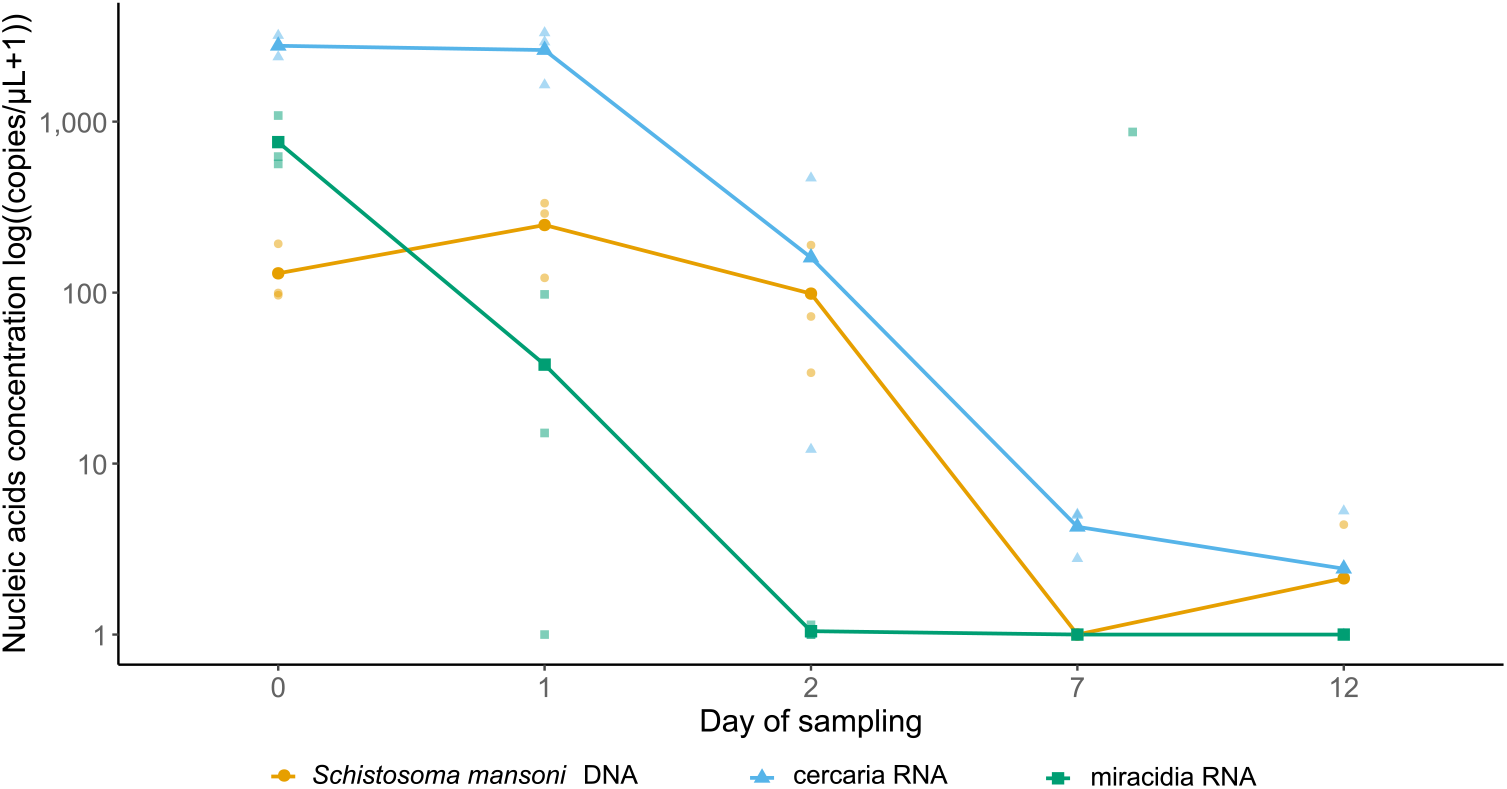
Temporal evolution of the concentrations in log((copies/µL+1)) of *S. mansoni* DNA (orange circle), cercariae RNA (blue triangle) and miracidia RNA (green square) for each kinetic point of the experiment (Day of sampling). Individual points represent sample measurements, lines connect the mean concentrations of nucleic acids among replicates at each kinetic point. Across all the experiment, negative-control tanks consistently remained free of detectable *S. mansoni* DNA and RNA.

Across all experiments, negative-control tanks consistently remained free of detectable *S. mansoni* DNA and RNA.

### Field validation

Field validation was conducted at six freshwater sites (ponds and streams) across three endemic localities (Dunga, Santo Antônio - in the city of Jaboticatubas, and Antônio Pereira – in Ouro Preto) in Minas Gerais state, Brazil. Classical parasitological surveys in *Biomphalaria glabrata* snails revealed the presence of *S. mansoni* at two sites, with infection prevalence of 5.4% at the site Dunga 1, and 0.8% at Antônio Pereira (Table 3). *Schistosoma mansoni* cercariae-specific eRNA, and to a lesser extent eDNA, were detected at the same two snail-positive sites (Table 3). At the remaining four sites where snails tested negative, a low concentration of cercariae-specific eRNA was detected in a single water replicate at Dunga 2 (out of 2 replicates), whereas eDNA remained undetected. No miracidia-specific eRNA signal was detected at any sites. Consistent with laboratory findings, stage-specific eRNA assays demonstrated higher sensitivity than eDNA assays in field samples (maximum concentrations: 64 copies/µL for cercariae-derived eRNA vs. 10.93 copies/µL for *S. mansoni* eDNA, Table 3). Also, using the average RNA copy number per 1 cercariae obtained in controlled conditions (SI Appendix, Text S1), we estimated that detected eRNA concentrations corresponded to approximately 0.04–0.2 cercariae-equivalent individuals, suggesting that signals primarily originated from released molecules rather than whole organisms, trapped during the water filtering process.

**Table 3.** Summary of *Schistosoma mansoni* prevalence in human at the municipality scale, and *Biomphalaria glabrata* abundance, *S. mansoni* prevalence in snail populations, average nucleic acids concentration (cercariae RNA, miracidia RNA, and Schistosoma mansoni DNA) at the sampling site scale. Field negative controls (Antônio Pereira - Ouro Preto) showed no detectable eRNA or eDNA (0 copies/µL), confirming the absence of contamination.

| Locality (City) | *S. mansoni*<br>prevalence in<br>human populations<br>at the municipality<br>scale (%) | Site | *Biomphalaria*<br>*glabrata*<br>abundance | *S. mansoni*<br>prevalence in snail<br>populations (%) | Cercariae<br>RNA<br>(copies/μL) | Miracidia<br>RNA<br>(copies/μL) | *S. mansoni*<br>DNA<br>(copies/μL) | Proportion of<br>positive filtration<br>replicate |
| --- | --- | --- | --- | --- | --- | --- | --- | --- |
| Dunga<br>(Jaboticatubas) | 28.21<br>(2025) | Dunga 1 | 223 | 5.4 | 14.73 | 0 | 1.23 | 2/2 |
|  |  | Dunga 2 | 6 | 0 | 0.13 | 0 | 0 | 1/2 |
|  |  | Dunga 3 | 0 | 0 | 0 | 0 | 0 | 0/2 |
| Santo Antônio<br>(Jaboticatubas) | 2.78<br>(2024) | Santo Antônio 1 | 57 | 0 | 0 | 0 | 0 | 0/2 |
|  |  | Santo Antônio 2 | 13 | 0 | 0 | 0 | 0 | 0/2 |
| Antônio Pereira<br>(Ouro Preto) | Not informed | Antônio Pereira | 131 | 0.8 | 64 | 0 | 10.93 | 1/2 |

## DISCUSSION

Here, we provide the first empirical evidence that environmental RNA can greatly improve the transmission surveillance of complex life-cycle parasites such as *S. mansoni* by enabling the detection of the infectious life stages that are actively circulating in the environment. We demonstrate the ability to specifically detect and quantify both cercariae and miracidia using eRNA-based approaches, hence providing a level of biological resolution that could not be previously achieved using eDNA for *S. mansoni* environmental surveillance. The specific detection of miracidia offers a direct, early, and non-invasive indicator of parasite input into freshwater environments by infected human or animal reservoir hosts and can therefore reveal aquatic sites under contamination pressure. When miracidia-specific signals co-occur with susceptible and compatible snail hosts, capable of subsequently amplifying and releasing cercariae, this can become an early-warning indicator of conditions permissive for upcoming transmission (14, 24). Miracidia detection from water samples can also uncover unrecognized infected human or animal host populations in the vicinity of the surveyed sites. The specific detection and quantification of cercariae-specific signal provides direct evidence of active ongoing infection risk, hence pinpointing active, infectious stages circulating in water bodies, which is an indicator with direct relevance for public-health decision-making. Beyond respective stage-specific detection, this approach enables investigation of the temporal dynamics of *S. mansoni* transmission. These include the seasonality of miracidia release by definitive hosts, cercariae shedding by snails, and resulting variation in human infection risk that are difficult to capture using traditional malacological or human surveys due to logistics and labor-intensity (25, 26). Consequently, this tool complements the existing toolbox for both better understanding the ecology of the transmission of *S. mansoni* and operationally managing transmission of intestinal schistosomiasis through the One Health approach. It opens new avenues for assessing the competence of aquatic ecosystems for *S. mansoni* as described by Douchet et al. (8), and for identifying environmental and ecological factors that enhance or suppress transmission at fine spatial and temporal scales.

In addition to providing life-stage resolution, our results demonstrate that stage-specific eRNA assays offer a greater sensitivity than eDNA for detecting *S. mansoni* from water samples. This challenges the prevailing assumption that DNA is inherently more stable and thus more readily detectable than RNA (22, 27, 28). Only a few pilot studies have demonstrated the feasibility of eRNA detection for some freshwater pathogens (29, 30), whereas here, both our experimental and field data align with recent studies reporting comparable or even superior detectability of eRNA relative to eDNA under certain conditions (21, 31). We consistently show that eRNA was more reliably detected than eDNA for both miracidia and cercariae. This pattern held true both when parasites were physically retained on filters and experimentally removed from water samples, indicating that eRNA assays also capture released free eRNA. This demonstrates that eRNA-based methods also effectively capture parasite signals from free environmental reservoirs, albeit at lower concentrations, which is expected in field conditions even for macrobial organisms (32). The higher sensitivity of eRNA detection likely results from the selection of highly expressed transcripts. This provides a natural amplification signal that can exceed the detectability of eDNA, including classical multi-copy mitochondrial targets used for *S. mansoni* detection (13, 15).

The degradation dynamics of eRNA revealed differential patterns between miracidia and cercariae. Despite a higher expected level of transcript expression, miracidia-derived eRNA signal rapidly declined and became undetectable at day 2, whereas cercariae-derived eRNA signal persisted up to days 7-12, exceeding the persistence of eDNA. This could lead to a lower detectability of miracidia compared to the cercaria signal in environmental and complex matrices. The longer persistence of cercariae-specific eRNA may be explained by cellular mechanisms such as transcript encapsulation within parasite-secreted extracellular vesicles (EVs). The cercariae transcript targeted here (*Smp_169190*), which encodes a tegumental allergen-like protein, may be protected from environmental degradation through this process. These vesicles, known to pack and protect mRNAs and microRNAs, shield RNA content against enzymatic degradation and environmental stress, thereby extending the temporal window of eRNA detection (33–36). Less is known about EV release and RNA cargo in miracidia, but their active secretory nature and reported RNA-rich vesicular output (33, 37) suggest that similar mechanisms may extend eRNA detectability for this stage as well, albeit to a lesser extent than in cercaria. Finally, while eRNA persistence may initially appear to limit eRNA’s capacity to distinguish living from dead organisms (20), it also enhances the robustness of detection in natural environments where parasite stages are highly transient, such as in *S. mansoni*. Environmental RNA persistence may therefore smooth short-term fluctuations linked to host behavior, parasite circadian rhythms, or seasonal variation, ultimately increasing detection reliability and improving long-term ecological surveillance of transmission sites (38, 39).

Our field survey in Minas Gerais, a region in Brazil characterized by low and heterogeneous schistosomiasis endemicity, validates the developed eRNA diagnostic tool for transmission surveillance. According to local health authorities, human prevalence of *S. mansoni* varied from 28.2% in Dunga locality to 2.78% in Santo Antônio locality, with no data available for Antônio Pereira, Ouro Preto. In this context, our eRNA assays, and to a lesser extent eDNA assays (which showed approximately six-fold lower concentrations), confirmed the presence of *S. mansoni* cercariae, indicative of ongoing transmission at one of the two human positive localities. It also provides novel epidemiological insights by detecting active cercariae in freshwater bodies of Ouro Preto, signaling ongoing transmission at a site so far lacking reported human cases. Moreover, environmental nucleic acids (eNAs) detection was consistent with the snail parasitological survey we simultaneously did at these six sites. Also, the eRNA concentrations measured at the two positive sites corresponded to RNA quantities less than those obtained from single cercariae, which suggests that we detected free or released eRNA and not whole cercariae. This likely indicates that sampling did not occur precisely at cercariae emission points. Deploying this method along spatial transects within water bodies could help identify cercariae emission hotspots. This would allow delineation of their associated microhabitats. Such spatially resolved surveillance would enhance targeting control interventions and environmental management efforts, crucial components of the WHO schistosomiasis control strategy, which emphasizes precision mapping and focal interventions to interrupt transmission effectively. Overall, these results demonstrate that stage-specific eRNA assays can be implemented under field conditions and provide robust epidemiological information beyond that obtained with conventional eDNA-based approaches.

Miracidia were not detected in field samples, suggesting no recent contamination of the water bodies studied by infected human or animal hosts. This finding aligns with our laboratory experiment, which showed that, even at high abundance, miracidia-derived eRNA signals are less readily detectable than cercariae and, while more detectable than eDNA the day of contamination, persist only briefly in the aquatic environment (< 2 days). This is also coherent with the well-established *S. mansoni* life cycle dynamics: miracidia, the short-lived larval stage emerging from eggs deposited by infected hosts, are shed intermittently and survive only briefly while seeking snail hosts, making their environmental detection challenging without intensive, longitudinal sampling. In contrast, infected snails release cercariae over days to weeks, enhancing their environmental persistence and detection probability (3, 40). Thus, cercariae detection confirms active transmission sites and human infection risk, whereas the absence of miracidia indicates no recent water contamination from infected definitive hosts, human or animal very locally and at the time of water sampling. Applying these approaches with finer spatial and temporal resolution at a local scale would allow more precise identification of specific sites along water bodies and the timing of parasite release by definitive hosts, whether human or animal.

While our method shows strong promise, improving field operationality as for eDNA protocols is the next step (11). Plankton nets, initially used in this study to concentrate *S. mansoni* nucleic acids and potential life stages, are labor-intensive and simpler direct filtration should be evaluated. Using slightly larger pore membranes (≈2.5–2.8 μm) could also boost recovery and detection of *S. mansoni* eDNA and eRNA (14). Finally, although no contamination was detected in our study, employing sealed membranes could further minimize risk, simplify handling, and enable safer, more efficient sample transport. Streamlining these steps will make stage-specific environmental RNA surveillance of schistosomes even more practical and scalable for routine monitoring.

More broadly, this proof-of-concept empirical study highlights the need to develop RNA-sensing technologies for *S. mansoni*, similar to those pioneered for viruses (41). Detecting stage-specific RNA transcripts for *S. mansoni* offers critical epidemiological insights into transmission dynamics of these parasites and the control of schistosomiasis. This improvement is particularly valuable for transmission surveillance in settings of low endemicity or to evaluate the efficiency of the eradication measured as in regions where human prevalence has declined following mass praziquantel administration hence aligning with WHO goals to eliminate schistosomiasis as a public health problem and interrupt transmission by 2030 (6). This study also calls for the development of such eRNA-based tools for other human-infecting schistosome species (e.g. *S. haematobium, S. japonicum*), which have broad geographic distributions across Africa and Asia, as well as to other parasites with complex life cycles and different infectious life stages of medical and veterinary significance. Achieving these goals requires comprehensive transcriptomic data covering all life stages of these parasite species. Ongoing development of multi-omics datasets for non-model parasites thus remains essential to underpin such novel diagnostic and surveillance strategies. Once developed for multiple pathogens, such environmental diagnostic tools could enable simultaneous monitoring of several parasites from a single water sample. This would increase the efficiency and accuracy of tracking transmission dynamics and infection risk in human communities. Beyond pathogen surveillance, this study opens the door to the use of eRNA for biodiversity assessment, ecosystem monitoring, or environmental functional ecology by enabling the detection of biologically active organisms and providing access to gene expression.

## MATERIALS & METHODS

### Stage-specific primers and probes design

The interactive ‘SchistoXYZ’ web platform (23) was used to identify genes preferentially expressed in *S. mansoni* miracidia but not in cercariae and *vice-versa* (SI Appendix, Fig S1). Ten genes were initially selected based on their maximal expression levels and their apparent specificities to each life stage. The corresponding exons sequences were retrieved from the WormBase database (42–44). For each selected gene we designed the primers and probes using the Primer3Plus tool (45) following the guidelines and the design constraints supplied with the Bio-Rad QX200 ddPCR system. To test the *in-silico* specificity of the designed primers, we used the Primer-BLAST tool (46) and both the nt nucleotide and Refseq RNA databases (NCBI), without targeting any particular organism (the other parameters were set as default).

To evaluate the effectiveness of eRNA-based approaches compared to recently developed eDNA-based approaches, we adapted the use of trematode-specific primers developed by Douchet et al. (47) that targets the mitochondrial *16S* gene coupled with a probe specific to *S. mansoni* designed in this study (Table 1).

### Validation of the specificity and sensitivity of the designed assays

The specificity and sensitivity of life stage-specific eRNA assays were evaluated using digital droplet PCR (ddPCR) on whole-organism RNA extracts from *S. mansoni* miracidia and cercariae. Stage specificity was assessed by testing each assay against RNA extracted from the alternate *S. mansoni* life stage. Species specificity was assessed using RNA extracted from a batch of frozen freshly hatched miracidia and cercariae from non-target trematode species. For cercariae, RNA was extracted from *Schistosoma haematobium*, and from two unidentified non-schistosome trematode species emitted by wild snails (*Ecrobia vitrea* and *Physa acuta*) collected during previous field campaigns in Southern France. For miracidia, RNA was extracted from *Schistosoma rodhaini, Fasciola hepatica* and *Calicophoron daubnei*. RNA extractions were performed using the E.Z.N.A. HP Total RNA kit (Omega Bio-tek), following the manufacturer’s instructions, adding a DNase treatment step with 15 minutes incubation. Equi-concentrated amounts of RNA extracts from each sample were then pooled to generate stage-specific mock communities excluding *S. mansoni* RNA.

Assay sensitivity was evaluated using RNA extracted in quadruplicate from frozen samples containing respectively 1 individual of *S. mansoni* miracidia or cercariae, using the same protocol. Limits of detection (LOD) were determined by preparing serial 1:10 dilutions of these RNA extract down to a 1:10000 dilution. The evaluation of specificity and sensitivity (LOD) for the *S. mansoni* eDNA assay is described in SI Appendix, Text S1.

RNA copy numbers were quantified using ddPCR on a QX200 system (Bio-Rad). PCR reactions were performed using the One-Step RT-ddPCR Advanced Kit for Probes (Bio-Rad®) in a total volume of 22 μL, containing 2 μL of RNA template, 1.98 μL of each stage-specific forward and reverse primers (900 nM), 0.55 μL of each corresponding probe (250 nM), 5.5 μL of 1X Digital PCR Supermix, 2.2 μL of reverse transcriptase (20 U/μL), 1.1 μL of 300 mM DTT, and 2.18 μL of DNase/RNase-free water, following the manufacturer’s recommendations. Reaction mixes were emulsified using the QX200 Droplet Generator (Bio-Rad®) and amplified using the following thermal program: 25°C for 3 min, 45°C for 60 min (reverse transcription), 95°C for 10 min, followed by 40 cycles of 95°C for 30 s and 60°C for 1 min, with a final enzyme deactivation step at 98°C for 10 min. Droplets were then read using the QX200 Droplet Reader (Bio-Rad®) to obtain absolute RNA quantification. For specificity assessments, each RNA template was analyzed in triplicate; for sensitivity and LOD determination, four technical replicates were performed. Each ddPCR run included no-template negative controls and positive controls consisting of *S. mansoni* miracidia- and cercariae-derived RNA.

### Experimental design and molecular processing for Hypotheses 1-3

To evaluate the performance of the stage-specific eRNA assays for quantifying relative life stage abundance (Hypothesis 1), detecting nucleic acid released into the water independently of whole organisms (Hypothesis 2), and characterizing the persistence and decay of eRNA and eDNA over time (Hypothesis 3), we conducted three controlled tank experiments under standardized laboratory conditions.

Prior to all experiments, laboratory facilities, tanks, and equipment were decontaminated with 70% ethanol and 10% bleach and rinsed with DNase/RNase-free water. Experimental tanks were filled with 3 L of untreated borehole water routinely used for parasite and snail maintenance at the IHPE laboratory (Perpignan, France). Freshly hatched miracidia and cercariae from a single *Schistosoma mansoni* lineage (*SmBre*), maintained at IHPE, were collected immediately prior to experiments. All experimental treatments were performed in triplicate.

After each experiment, the entire water volume from each tank was filtered on a 0.45 µM polyethersulfone (PES) membrane mounted in a filter holder (eDNA Filter Pack, Smith-Root) using an electric pump powered by a 12-V battery. Each membrane was halved and placed into 1 mL of DNA/RNA Lysis Buffer (Zymo Research) and stored at room temperature for up to 72 h. Prior to extraction, tubes were agitated at 900 rpm for 2 h. Lysates from both filter halves were pooled and processed using the Quick-DNA/RNA MiniPrep Plus Kit (Zymo Research), allowing simultaneous DNA and RNA extraction, with a 25-min DNase-I treatment to eliminate genomic DNA from RNA extracts. Extractions were conducted according to the manufacturer’s recommendations. To maximize nucleic acid recovery elution was performed twice, yielding 50 μL of purified nucleic acids. RNA aliquots (2.2 μL) were stored at −70 °C and DNA extracts were stored at −20 °C.

Nucleic acid quantification was performed using ddPCR, as described above. RNA quantification used the One-Step RT-ddPCR Advanced Kit for Probes (Bio-Rad), and DNA quantification used the ddPCR Supermix for Probes (No dUTP, Bio-Rad). To assess potential residual DNA contamination, RNA extracts were also amplified using the DNA-specific Supermix. When detected, residual DNA copy numbers were subtracted from total RNA copy numbers. Each plate included negative controls (DNase/RNase-free water) and positive controls (extracted *S. mansoni* DNA, RNA from either cercariae or miracidia, and mock communities of miracidia or cercariae)

### Hypothesis 1 - Stage-specific eRNA reliably quantify *S. mansoni* life stages abundance

To test whether stage-specific eRNA accurately reflects the relative abundance of *S. mansoni* life stages and compare it to eDNA, we filtered water from tanks containing three ratios of *S. mansoni* cercariae and miracidia: balanced ratios (25:25) and imbalanced (5:45; 45:5). All freshly hatched *S. mansoni* free-living stages were first isolated in batches in 1.5 ml tubes and then released in their respective experimental tanks simultaneously. An additional tank with no *S. mansoni* was also prepared as a negative control. Water filtration occurred 5 hours after organism introduction.

Here, we set a Generalized Linear Model (GLM) setting the nucleic acid concentration (*i*.*e*. each eRNA specific transcripts and *S. mansoni* DNA) as a dependent variable and the nucleic acid assay type and the experimental ratio conditions (miracidia/cercariae: 5:45, 25:25, 45:5) as explanatory variables accounting for their potential interaction. We applied a Tukey-adjusted post-hoc contrasts analysis to assess pairwise differences among nucleic acid types and ratios. Based on the mean eRNA concentrations obtained for each life stage in the 25:25 ratio condition, we calculated the expected average eRNA concentrations for the 5:45 and 45:5 ratios and compared these expected values with the corresponding observed concentrations.

### Hypothesis 2 - Released eRNA and eDNA are detectable

To assess the detectability of released nucleic acids that are not associated with whole free-living organisms, we set up a two steps filtration experiment. Prior to water filtration, 5 hours after exposure, three tanks with balanced cercariae and miracidia ratios (25:25) were pre-filtered through a 25 µM mesh sieve to remove the entire organisms. The pre-filtered collected water was then directly filtered on 0.45 μm PES membrane. These results were then compared with triplicate tanks containing organisms which were processed identically.

Here, we expected to detect shed RNA and DNA, albeit at lower concentrations than in treatments containing live organisms. A negative-binomial model was set to test the effects of the nucleic acid type and the experimental treatment (i.e. organisms removed vs. present), accounting for an interaction term. We also applied a Tukey-adjusted contrasts among experimental treatment and among nucleic acid assay type.

### Hypothesis 3 - eRNA persists as long as eDNA signal in experimental tanks

To characterize the temporal persistence and decay of eRNA and eDNA, 15 water tanks were seeded with balanced proportion of miracidia and cercariae on day 0 (25:25), and five additional tanks served as *S. mansoni* free controls. Water was then filtered at five kinetic points: day 0 (after 5 hours), day 1, day 2, day 7 and day 12. Because miracidia and cercariae generally live up to 24 h (48), dead organisms were left in tanks to approximate natural degradation processes. Three replicated tanks plus one negative control was processed per kinetic point.

A negative binomial model was set to test the effects of the nucleic acid assay type, and the sampling day including an interaction term on nucleic acid concentration which was set as the dependent variable. Post-hoc contrasts compared temporal decay trajectories among each nucleic acid assay type (cercariae RNA, miracidia RNA and *S. mansoni* DNA).

All the statistical analyses were performed using R version 4.4.1 (R Core Team, 2024) using the glm.nb function implemented in the *MASS* package and the Tukey-adjusted post-hoc contrasts were done using the *emmeans* package .

### Field validation

Field validation of the eRNA -based environmental surveillance tool was conducted at six freshwater sites (ponds and streams) across three schistosomiasis-endemic localities—Dunga, Santo Antônio, and Ouro Preto—in Minas Gerais state, Brazil, in May 2025. Epidemiological data on *S. mansoni* prevalence human populations were available for Dunga and Santo Antônio but not for Ouro Preto locality. Each site investigated were located in endemic areas, hosted established populations of the intermediate snail host *Biomphalaria glabrata*, and were accessible within three hours by car from the Fiocruz laboratory.

At each site, water samples were collected prior to snail sampling, and physicochemical parameters (T°C, pH, conductivity, hardness) were recorded (SI Appendix, Table S7). Snail collection was assessed along the same water bodies as for water sampling, using metal scoops for 15 minutes by two operators per each site. Snails were stored alive in labelled plastic bags, and transported to the laboratory to test for cercariae emission. Snails were individually tested for cercaria shedding using light exposure for 24 h, and emitted cercariae were examined under microscope to identify the presence of *S. mansoni* cercariae and calculate the prevalence (50).

Environmental nucleic acids samples were collected in duplicates using a plankton net (25 cm diameter, 30 µm mesh) pulled along a 10 m transect, concentrating approximately 490 L of filtered water into a 500 mL collection bottle (placed at the net’s exit). Each 500 mL sample was filtered through PES membranes (0.45 μm), which were replaced when saturated. Filters were cut in half and preserved in 1 mL of DNA/RNA Lysis Buffer (Zymo Research) within sterile 1.5 mL tubes, then stored at 4°C until subsequent molecular processing.

To prevent cross-contamination, a dedicated set of sampling equipment (net, collection bottle, and filtration holder) was used per site and decontaminated between replicates using 10% bleach followed by thorough rinsing with clean water. Field processing of membranes included successive decontamination baths in ethanol, DNA Away, and Milli-Q water, with sterile gloves changed between each water replicate. One field negative control was done at Ouro Preto by passing drinking water through the net, filtered in the 500 mL bottle.

RNA and DNA extractions followed previous experimental protocols used, with the use of two purification columns per sample to prevent membrane saturation. Eluates from both columns per sample were pooled prior to downstream analyses. Quantification of *S. mansoni* eDNA and stage-specific eRNA was performed using the QX200 AutoDG Droplet Digital PCR System (Bio-Rad®). Negative controls were included at each molecular step, and positive controls from cercariae and miracidia-derived RNA were included in all ddPCR runs.

## Acknowledgments

We thank Julie Clement (IHPE) and Kleiton Esteves Costa and Fellipe Simasat from Fiocruz René Rachou for their help and assistance during field work validation; the disease control agents from the state of Minas Gerais, municipalities of Jaboticatubas and Ouro Preto for their technical support in the fieldwork; Ronaldo De Carvalho Augusto (IHPE), Bertrand de l’Isle (IHPE) and Emeline Félix (S.A.S. ParaDev) for maintaining and providing living schistosomes in laboratory. The ddPCRs were run thanks to the Bio-Environment platform (University of Perpignan Via Domitia, Occitanie Region) with the assistance of Margot Doberva. CM, OR, SB, and GL were supported by the ALIQUOT project (ANR-22-PEXO-0011). Funding from the EUR TULIP-GSR N°ANR-18-EUR-0019 was granted to OR and CM. OR was also supported by the Occitanie Region (Schistodiag program). This study is set within the framework of the « Laboratoire d’Excellence (LabEx) » TULIP (ANR-10-LABX-41).

## Notes

### Competing Interest Statement

The authors have declared no competing interest.

