## Supplementary Information for "Environmental RNA Improves Detection and Surveillance of Schistosomiasis Transmission"

**Figure S1:**

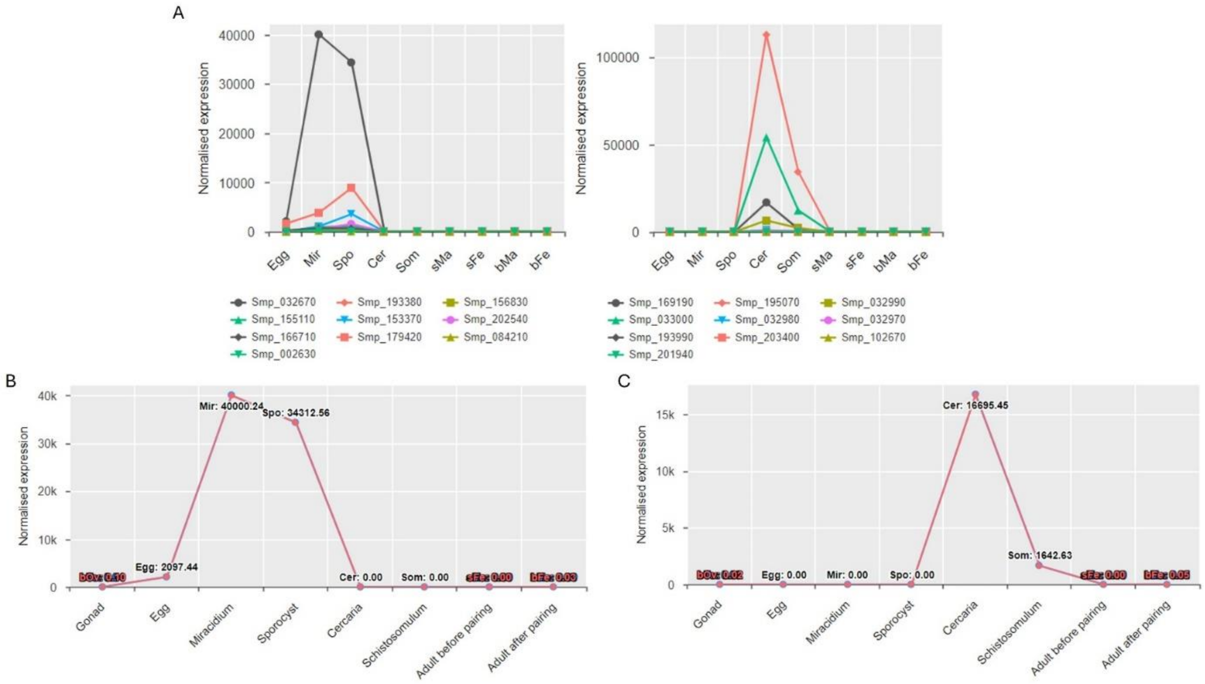


**Fig. S1. Gene expression profiles across *S. mansoni* life stages from the 'SchistoXYZ' platform. (A) left panel: expression profile across each life stages of the top 10 genes specifically expressed in miracidia and sporocyst; right panel: expression profile across each life stages of the top ten genes specifically expressed in cercariae and schistosomula. (B) expression profile across each life stages of Smp_032670, targeted gene for miracidia. (C) expression profile across each life stages of Smp_169190, targeted gene for cercaria.**

### Figure S2:

| | Sequence (5'->3') | Length | Tm | GC% | Self complementarity | Self 3' complementarity |
|---|---|---|---|---|---|---|
| **Forward primer** | CTGAACAGACTGCCGTGGTG | 20 | 61.22 | 60.00 | 5.00 | 2.00 |
| **Reverse primer** | GTGTCGCTAATCACTTCTGAGC | 22 | 59.40 | 50.00 | 3.00 | 3.00 |

**Products on target templates**

>XM_018795249.1 Schistosoma mansoni egg protein C122-like partial mRNA

```
product length = 106
Forward primer  1     CTGAACAGACTGCCGTGGTG  20
Template        172   ....................  191

Reverse primer  1     GTGTCGCTAATCACTTCTGAGC  22
Template        277   ......................  256
```

>XM_018795250.1 Schistosoma mansoni egg protein C122-like partial mRNA

```
product length = 106
Forward primer  1     CTGAACAGACTGCCGTGGTG  20
Template        137   ....................  156

Reverse primer  1     GTGTCGCTAATCACTTCTGAGC  22
Template        242   ......................  221
```

>HE601625.3 Schistosoma mansoni strain Puerto Rico genome assembly, chromosome: 2

```
product length = 106
Forward primer  1          CTGAACAGACTGCCGTGGTG  20
Template        43789939   ....................  43789920

Reverse primer  1          GTGTCGCTAATCACTTCTGAGC  22
Template        43789834   ......................  43789855
```

**Fig. S2 a. Primer Blast results for the miracidia-Primers on nt database.**

| | Sequence (5'->3') | Length | Tm | GC% | Self complementarity | Self 3' complementarity |
|---|---|---|---|---|---|---|
| **Forward primer** | CTGAACAGACTGCCGTGGTG | 20 | 61.22 | 60.00 | 5.00 | 2.00 |
| **Reverse primer** | GTGTCGCTAATCACTTCTGAGC | 22 | 59.40 | 50.00 | 3.00 | 3.00 |

**Products on target templates**

>XM_018795249.1 Schistosoma mansoni egg protein C122-like partial mRNA

```
product length = 106
Forward primer  1     CTGAACAGACTGCCGTGGTG  20
Template        172   ....................  191

Reverse primer  1     GTGTCGCTAATCACTTCTGAGC  22
Template        277   ......................  256
```

>XM_018795250.1 Schistosoma mansoni egg protein C122-like partial mRNA

```
product length = 106
Forward primer  1     CTGAACAGACTGCCGTGGTG  20
Template        137   ....................  156

Reverse primer  1     GTGTCGCTAATCACTTCTGAGC  22
Template        242   ......................  221
```

**Fig. S2 b. Primer Blast results for the miracidia-Primers on RefSeq RNA database.**

| | Sequence (5'->3') | Length | Tm | GC% | Self complementarity | Self 3' complementarity |
|---|---|---|---|---|---|---|
| **Forward primer** | GTACGGAATACCTACAACGG | 20 | 55.41 | 50.00 | 6.00 | 1.00 |
| **Reverse primer** | ATGTCGTGTCAAGCCATC | 18 | 55.04 | 50.00 | 7.00 | 3.00 |

**Products on target templates**

>OX104036.1 Schistosoma rodhaini genome assembly, chromosome: 1

```
product length = 173
Forward primer  1          GTACGGAATACCTACAACGG  20
Template        18764920   ....................  18764901

Reverse primer  1          ATGTCGTGTCAAGCCATC  18
Template        18764748   ..................  18764765
```

>XM_018794911.1 Schistosoma mansoni putative tegumental protein partial mRNA

```
product length = 173
Forward primer  1     GTACGGAATACCTACAACGG  20
Template        283   ....................  302

Reverse primer  1     ATGTCGTGTCAAGCCATC  18
Template        455   ..................  438
```

>OX103911.1 Schistosoma rodhaini genome assembly, chromosome: 1

```
product length = 173
Forward primer  1          GTACGGAATACCTACAACGG  20
Template        68576597   ....................  68576616

Reverse primer  1          ATGTCGTGTCAAGCCATC  18
Template        68576769   ..................  68576752
```

>HE601624.3 Schistosoma mansoni strain Puerto Rico genome assembly, chromosome: 1

```
product length = 173
Forward primer  1          GTACGGAATACCTACAACGG  20
Template        70012937   ....................  70012956

Reverse primer  1          ATGTCGTGTCAAGCCATC  18
Template        70013109   ..................  70013092
```

**Fig. S2 c. Primer Blast results for the cercariae-Primers on nt database.**

| | Sequence (5'->3') | Length | Tm | GC% | Self complementarity | Self 3' complementarity |
|---|---|---|---|---|---|---|
| **Forward primer** | GTACGGAATACCTACAACGG | 20 | 55.41 | 50.00 | 6.00 | 1.00 |
| **Reverse primer** | ATGTCGTGTCAAGCCATC | 18 | 55.04 | 50.00 | 7.00 | 3.00 |

**Products on target templates**

>XM_018794911.1 Schistosoma mansoni putative tegumental protein partial mRNA

```
product length = 173
Forward primer  1    GTACGGAATACCTACAACGG  20
Template        283  ....................  302

Reverse primer  1    ATGTCGTGTCAAGCCATC  18
Template        455  ..................  438
```

>XM_012941916.1 Schistosoma haematobium Tegument antigen (A12_9), partial mRNA

```
product length = 173
Forward primer  1    GTACGGAATACCTACAACGG  20
Template        231  ..................A.  250

Reverse primer  1    ATGTCGTGTCAAGCCATC  18
Template        403  ....T.............  386
```

**Fig. S2 d. Primer Blast results for the cercariae-Primers on RefSeq RNA database.**

**Table S1: Limit of Detection (LOD) measurement for eRNA assays.***

**Table S1 a. LOD for RNA stage specific primers and probe to target miracidia.**

| Dilution of 1 individual | Life stage sampled | Mean RNA concentration (copies/µL) | % detected replicates |
|---|---|---|---|
| 0.1 | Miracidia | 8.52 ± 4.29 | 100% |
| 0.01 | Miracidia | 0.73 ± 0.72 | 100% |
| 0.001 | Miracidia | 0.0186 ± 0.0373 | 25% |

**Table S1 b. LOD for RNA stage specific primers and probe to target cercariae.**

| Dilution of 1 individual | Life stage sampled | Mean RNA concentration (copies/µL) | % detected replicates |
|---|---|---|---|
| 0.01 | Cercaria | 5.01 ± 4.45 | 100% |
| 0.001 | Cercaria | 0.72 ± 0.43 | 100% |
| 0.0001 | Cercaria | 0.139 ± 0.053 | 100% |

*To determine the LOD of the cercariae and miracidia specific primers we made serial dilution 1:10 up to 1/10.000 from the 1 miracidia and 1 cercaria individual in quadruplicate. We considered the LOD as the lowest dilution for which 100% of the samples are positive, and then quantifiable by dPCR. According to the concentrations obtained for each replicate, the LOD of the RNA designed primers are 0.01 individual for miracidia and 0.0001 individual for cercariae

**Text S1: Methods and results for the specificity and sensitivity evaluation of the *16S* eDNA assay for *Schistosoma mansoni* quantification**.

The specificity of the *16S* eDNA assay targeting *S. mansoni* was tested using equimolar mock community constructed with extracted DNA of 5 trematode species including two different strains of *Dicrocoelium dendriticum*, two different strains of *Schistosoma haematobium*, *Fasciola hepatica*, *Fasciola gigantica* and *Posthodiplostomum centrarchi*. The LOD of the *S. mansoni* eDNA assay was measured by serial 1:10 dilutions prepared down to 1/1000 from DNA respectively extracts of a single miracidia and a single cercaria in quadruplicate. DNA extraction was made with the Quick-DNA/RNA MiniPrep Plus kit (Zymo Research). The DNA detection and quantification were performed based on ddPCR assays using a Bio-Rad QX200 system. PCR reactions were performed using the ddPCR Supermix for Probes (No dUTP, Bio-Rad®) in a total reaction volume of 22 µL, following the manufacturer's recommendations on concentrations. Individual reaction mix were emulsified using the QX200 Droplet Generator (Bio-Rad®) and amplified using the following PCR program: 25°C for three min, 95°C for 10 min, 40 cycles of 30 sec at 95°C followed by one min at 60°C and 98°C for 10 min. Absolute quantification of *Schistosoma mansoni* DNA was subsequently performed using the QX200 Droplet Reader (Bio-Rad®).

The specificity test performed with mock community confirmed the specificity of the eDNA designed primers and probe to *S. mansoni.* No amplification was observed in the sample without the presence of *Schistosoma mansoni* (Table 1a). The LOD of the DNA designed probe was 0.01 individual no matter the life stages (Table 1b).

**Table 1a. *S. mansoni* DNA concentration (in copies/µL) obtained in each mock community constructed for testing specificity of the eDNA assays using ddPCR.**

| Sample tested | Raw *S. mansoni* DNA concentration (copies/µL) |
|---|---|
| Mock community with *S. mansoni* | 10.1 |
| Mock community without *S. mansoni* | 0 |

**Table 1b. LOD results for Schistosoma mansoni DNA assays.**

| Dilution of 1 individual | Life stage sampled | Mean DNA concentration (copies/µL) | % detected replicates |
|---|---|---|---|
| 0.1 | Cercariae | 1.44 ± 0.42 | 100% |
| 0.01 | Cercariae | 0.2 ± 0.10 | 100% |
| 0.001 | Cercariae | 0.02 ± 0.03 | 25% |
| 0.1 | Miracidia | 1.88 ± 0.47 | 100% |
| 0.01 | Miracidia | 0.25 ± 0.23 | 100% |
| 0.001 | Miracidia | 0.01 ± 0.031 | 25% |

**Table S2: Parameter estimates obtained from the GLM model explaining the nucleic acid concentration in the conditions of the Hypothesis 1: eRNA reliably quantify *S. mansoni* life stages.***

| Variable | Deviance | Residual df | Residual deviance | p-value |
|---|---|---|---|---|
| Nucleic acid assay type | 268.62 | 21 | 141.16 | < 0.001 |
| Ratio condition | 9.18 | 19 | 131.98 | < 0.01 |
| Nucleic acid assay type*Ratio condition | 107.07 | 15 | 24.92 | < 0.001 |

* The GLM model was constructed with a Negative Binomial distribution as follow: nucleic acid concentration ~ Nucleic acid type*Ratio condition. The nucleic acid type considered 3 options: cercariae RNA, miracidia RNA and *S. mansoni* DNA. The ratio conditions considered 3 options: 45:5, 25:25, 5:45.

**Table S3: Post hoc pairwise contrasts tests for nucleic acid concentration across miracidia:cercariae ratios (a) among the proportion condition, and (b) among the nucleic acid type in the test of the Hypothesis 1.**

| (a) | Estimate | SE | z-ratio | p-value |
|---|---|---|---|---|
| **Ratio 45:5** | | | | |
| DNA *S. mansoni* - RNA cercariae | -0.903 | 0.318 | -2.840 | 0.0125 |
| DNA *S. mansoni* - RNA miracidia | -2.029 | 0.317 | -6.403 | <.001 |
| RNA S. cercariae - RNA miracidia | -1.126 | 0.315 | -3.576 | 0.001 |
| **Ratio 25:25** | | | | |
| DNA *S. mansoni* - RNA cercariae | -3.072 | 0.261 | -11.769 | <.0001 |
| DNA *S. mansoni* - RNA miracidia | -1.773 | 0.262 | -6.779 | <.0001 |
| RNA S. cercariae - RNA miracidia | 1.298 | 0.257 | 5.055 | <.0001 |
| **Ratio 5:45** | | | | |
| DNA *S. mansoni* - RNA cercariae | -3.316 | 0.260 | -12.759 | <.0001 |
| DNA *S. mansoni* - RNA miracidia | 0.119 | 0.264 | 0.450 | 0.8945 |
| RNA S. cercariae - RNA miracidia | 3.435 | 0.260 | 13.191 | <.0001 |

| (b) | Estimate | SE | z-ratio | p-value |
|---|---|---|---|---|
| ***S. mansoni* DNA concentration** | | | | |
| 45:5 – 25:25 | 0.641 | 0.294 | 2.181 | 0.0745 |
| 45:5 – 5:45 | 0.425 | 0.293 | 1.449 | 0.3156 |
| 25:25 – 5:45 | -0.216 | 0.265 | -0.817 | 0.6927 |
| **Cercariae RNA concentration** | | | | |
| 45:5 – 25:25 | -1.528 | 0.288 | -5.314 | <.0001 |
| 45:5 – 5:45 | -1.989 | 0.288 | -6.919 | <.0001 |
| 25:25 – 5:45 | -0.461 | 0.256 | -1.800 | 0.1695 |
| **Miracidia RNA concentration** | | | | |
| 45:5 – 25:25 | 0.897 | 0.287 | 3.122 | 0.0051 |
| 45:5 – 5:45 | 2.573 | 0.290 | 8.857 | <.0001 |
| 25:25 – 5:45 | 1.676 | 0.261 | 6.418 | <.0001 |

**Table S4: Parameter estimates obtained from the GLM model explaining the nucleic acid concentration in the conditions of the Hypothesis 2: True eRNA and eDNA signals can be detectable. ***

| **Variable** | **Deviance** | **Residual Df** | **Residual deviance** | **Pr(>Chi)** |
|---|---|---|---|---|
| Experimental treatment | 107.315 | 16 | 53.140 | < 0.0001 |
| Nucleic acid assay type | 28.299 | 14 | 24.841 | < 0.0001 |
| Experimental treatment*Nucleic acid type | 5.328 | 12 | 19.513 | 0.07 |

* The GLM model was constructed as follow: Nucleic acid concentration ~ Nucleic acid assay type*Experimental treatment. The nucleic acid assay type considered 3 options: cercaria RNA, miracidia RNA and *S. mansoni* DNA. The experimental treatment considered 2 options: with organisms and without organisms.

**Table S5: Parameter estimates obtained from the GLM model explaining the nucleic acid concentration in the conditions of the Hypothesis 3: eRNA signal persists as long as eDNA signal in experimental tanks.***

| **Variable** | **Deviance** | **Residual Df** | **Residual deviance** | **Pr(>Chi)** |
|---|---|---|---|---|
| Day of sampling | 318.07 | 40 | 135.33 | < 0.001 |
| Nucleic acid assay type | 47.79 | 38 | 87.54 | < 0.001 |
| Day of sampling*Nucleic acid type | 47.59 | 30 | 39.95 | < 0.001 |

* The GLM model was constructed with a Negative Binomial distribution as follow: Nucleic acid concentration ~ Nucleic acid assay type*Day of sampling. The nucleic acid assay type considered 3 options: cercariae RNA, miracidia RNA and *S. mansoni* DNA. The day of sampling considered 3 options: day 0, day 1, day 2, day 7, day 12.

**Table S6: Post hoc pairwise contrasts for nucleic acid concentration temporal persistence by days among the among the nucleic acid type in the Hypothesis 3.**

| | Estimate | SE | z.ratio | p.value |
|---|---|---|---|---|
| ***S. mansoni* DNA concentration** | -0.6505 | 0.621 | -1.048 | 0.8330 |
| Day_of_sampling0 - Day_of_sampling1 | -0.6505 | 0.621 | -1.048 | 0.8330 |
| Day_of_sampling0 - Day_of_sampling2 | 0.2732 | 0.622 | 0.439 | 0.9923 |
| Day_of_sampling0 - Day_of_sampling7 | 4.8650 | 0.847 | 5.744 | <.0001 |
| Day_of_sampling0 - Day_of_sampling12 | 4.1081 | 0.735 | 5.588 | <.0001 |
| Day_of_sampling1 - Day_of_sampling2 | 0.9237 | 0.621 | 1.486 | 0.5714 |
| Day_of_sampling1 - Day_of_sampling7 | 5.5154 | 0.846 | 6.517 | <.0001 |
| Day_of_sampling1 - Day_of_sampling12 | 4.7585 | 0.734 | 6.480 | <.0001 |
| Day_of_sampling2 - Day_of_sampling7 | 4.5917 | 0.847 | 5.418 | <.0001 |
| Day_of_sampling2 - Day_of_sampling12 | 3.8348 | 0.736 | 5.213 | <.0001 |
| Day_of_sampling7 - Day_of_sampling12 | -0.7569 | 0.933 | -0.811 | 0.9274 |
| **Cercariae RNA concentration** | | | | |
| Day_of_sampling0 - Day_of_sampling1 | 0.0573 | 0.618 | 0.093 | 1.0000 |
| Day_of_sampling0 - Day_of_sampling2 | 2.8527 | 0.619 | 4.605 | <.0001 |
| Day_of_sampling0 - Day_of_sampling7 | 6.4780 | 0.678 | 9.555 | <.0001 |
| Day_of_sampling0 - Day_of_sampling12 | 7.0427 | 0.720 | 9.776 | <.0001 |
| Day_of_sampling1 - Day_of_sampling2 | 2.7953 | 0.619 | 4.513 | 0.0001 |
| Day_of_sampling1 - Day_of_sampling7 | 6.4206 | 0.678 | 9.470 | <.0001 |
| Day_of_sampling1 - Day_of_sampling12 | 6.9854 | 0.720 | 9.697 | <.0001 |
| Day_of_sampling2 - Day_of_sampling7 | 3.6253 | 0.679 | 5.336 | <.0001 |
| Day_of_sampling2 - Day_of_sampling12 | 4.1900 | 0.722 | 5.805 | <.0001 |
| Day_of_sampling7 - Day_of_sampling12 | 0.5647 | 0.773 | 0.731 | 0.9494 |
| **Miracidia RNA concentration** | | | | |
| Day_of_sampling0 - Day_of_sampling1 | 2.9965 | 0.625 | 4.794 | <.0001 |
| Day_of_sampling0 - Day_of_sampling2 | 6.5863 | 0.837 | 7.870 | <.0001 |
| Day_of_sampling0 - Day_of_sampling7 | 6.6319 | 0.846 | 7.842 | <.0001 |
| Day_of_sampling0 - Day_of_sampling12 | 6.6319 | 0.846 | 7.842 | <.0001 |
| Day_of_sampling1 - Day_of_sampling2 | 3.5898 | 0.842 | 4.264 | 0.0002 |
| Day_of_sampling1 - Day_of_sampling7 | 3.6354 | 0.851 | 4.274 | 0.0002 |
| Day_of_sampling1 - Day_of_sampling12 | 3.6354 | 0.851 | 4.274 | 0.0002 |
| Day_of_sampling2 - Day_of_sampling7 | 0.0456 | 1.020 | 0.045 | 1.0000 |
| Day_of_sampling2 - Day_of_sampling12 | 0.0456 | 1.020 | 0.045 | 1.0000 |
| Day_of_sampling7 - Day_of_sampling12 | 0.0000 | 1.020 | 0.000 | 1.0000 |

**Table S7: Sites description and abiotic parameters measured at the water and snail collection sites.**

| Sites | Type of freshwater | Geographical coordonates | Conductivity (µS/cm) | Hardness (calcium) (mg/L) | pH | Water temperature (°C) |
|---|---|---|---|---|---|---|
| **Dunga 1** | Stream | 19°34’59” S 43°46’48” W | 171 | 130 | 7.5 | 22.4 |
| **Dunga 2** | Pond | 19°35’32” S 43°46’53” W | 215.6 | 130 | 7.9 | 22.24 |
| **Dunga 3** | Stream | 19°33’55” S 43°46’46” W | 266 | 125 | 7.7 | 22.6 |
| **Santo Antonio 1** | River | 19°24’35” S 43°42’33” W | 121.7 | 27,6 | 6.9 | 20.6 |
| **Santo Antonio 2** | River | 19°24’38” S 43°42’30” W | 78 | 24 | 7.13 | 19.26 |
| **Antônio Pereira** | River | 20°18’14” S 43°29’04” w | 53.3 | 38.3 | 7.5 | 24.6 |
